## Supplementary Information for "Efficient computation of adjoint sensitivities at steady-state in ODE models of biochemical reaction networks"

### Contents

**S1 Adjoint state integral computation at steady state** **S2**

**S2 Conversion reaction example** **S2**

**S3 AMICI implementation** **S4**

**S4 Simulation during optimization with ssASA takes less time** **S5**

### S1 Adjoint state integral computation at steady state

We assume that the system

$$\dot{\mathbf{x}} = \mathbf{f}(\mathbf{x}(t, \boldsymbol{\theta}, \mathbf{u}), \boldsymbol{\theta}, \mathbf{u}),\ \mathbf{x}(t_0, \boldsymbol{\theta}) = \mathbf{x}_0(\boldsymbol{\theta}, \mathbf{u}), \tag{1}$$

has an exponentially stable steady state, which means that the real parts of the eigenvalues of the Jacobian at the steady state ($\mathbf{J}(\mathbf{x}^*(\boldsymbol{\theta}, \mathbf{u}), \boldsymbol{\theta}, \mathbf{u})$) are negative. The system

$$\dot{\mathbf{p}}(t, \boldsymbol{\theta}, \mathbf{u}) = -\mathbf{J}(\mathbf{x}^*(\boldsymbol{\theta}, \mathbf{u}), \boldsymbol{\theta}, \mathbf{u})^T \mathbf{p}(t, \boldsymbol{\theta}, \mathbf{u}) \tag{2}$$

has a steady state $\mathbf{p} = \mathbf{0}$, which is then, as the real parts of the eigenvalues of $\mathbf{J}(\mathbf{x}^*(\boldsymbol{\theta}, \mathbf{u}), \boldsymbol{\theta}, \mathbf{u})$ are negative, asymptotically stable in reverse time. Hence, on the interval $[t', t'')$, where the system (1) is at steady state,

$$\begin{aligned}
&\mathbf{p}_{\text{integral}} \\
&= \int_{t'}^{t''} \mathbf{p}(s, \boldsymbol{\theta}, \mathbf{u}) ds \\
&= \int_{t'}^{t''} e^{-\mathbf{J}(\mathbf{x}^*(\boldsymbol{\theta}, \mathbf{u}), \boldsymbol{\theta}, \mathbf{u})^T (s - t'')} \mathbf{p}(t'') ds \\
&= -\left(\mathbf{J}(\mathbf{x}^*(\boldsymbol{\theta}, \mathbf{u}), \boldsymbol{\theta}, \mathbf{u})^T\right)^{-1} \mathbf{p}(t'') e^{-\mathbf{J}(\mathbf{x}^*(\boldsymbol{\theta}, \mathbf{u}), \boldsymbol{\theta}, \mathbf{u})^T (t'' - t'')} \\
&\qquad + \underbrace{\left(\mathbf{J}(\mathbf{x}^*(\boldsymbol{\theta}, \mathbf{u}), \boldsymbol{\theta}, \mathbf{u})^T\right)^{-1} \mathbf{p}(t'') e^{-\mathbf{J}(\mathbf{x}^*(\boldsymbol{\theta}, \mathbf{u}), \boldsymbol{\theta})^T (t' - t'')}}_{=0} \\
&= -\left(\mathbf{J}(\mathbf{x}^*(\boldsymbol{\theta}, \mathbf{u}), \boldsymbol{\theta}, \mathbf{u})^T\right)^{-1} \mathbf{p}(t''),
\end{aligned}$$

as

$$e^{-\mathbf{J}(\mathbf{x}^*(\boldsymbol{\theta}, \mathbf{u}), \boldsymbol{\theta}, \mathbf{u})^T (t' - t'')} = 0.$$

The computed $p_{\text{integral}}$ value is used to compute the objective function gradient by

$$\begin{aligned}
\left.\frac{\partial \mathcal{J}}{\partial \theta_k}\right|_{\boldsymbol{\theta}} = &-\sum_{i=1}^{n_y} \sum_{j=1}^{n_t} \frac{(\bar{y}_{ij} - y_i(t_j, \boldsymbol{\theta}, \mathbf{u}))}{\sigma_{ij}^2} \left.\frac{\partial h_i}{\partial \theta_k}\right|_{\mathbf{x}(t, \boldsymbol{\theta}, \mathbf{u}), \boldsymbol{\theta}, \mathbf{u}} - \sum_{i=1}^{n_y} \frac{(\bar{y}_{i*} - y_i(t'', \boldsymbol{\theta}, \mathbf{u}))}{\sigma_{i*}^2} \left.\frac{\partial h_i}{\partial \theta_k}\right|_{\mathbf{x}(t, \boldsymbol{\theta}, \mathbf{u}), \boldsymbol{\theta}, \mathbf{u}} \\
&- \int_{t_0}^{t''} \mathbf{p}(t, \boldsymbol{\theta}, \mathbf{u})^T \left.\frac{\partial \mathbf{f}}{\partial \theta_k}\right|_{\mathbf{x}(t, \boldsymbol{\theta}, \mathbf{u}), \boldsymbol{\theta}, \mathbf{u}} dt - \mathbf{p}(t_0, \boldsymbol{\theta}, \mathbf{u})^T \left.\frac{\partial \mathbf{x}_0}{\partial \theta_k}\right|_{\boldsymbol{\theta}, \mathbf{u}}.
\end{aligned}$$

### S2 Conversion reaction example

In this section, we illustrate the proposed method on an exemplary conversion reaction

$$\mathrm{A} \underset{\theta_2}{\overset{\theta_1}{\rightleftharpoons}} \mathrm{B},$$

where reaction rate coefficients $\theta_1, \theta_2 > 0$. We perform the computations from Section "Methods, Adjoint sensitivity analysis at steady state, Post-equilibration case" for this simple case.

The ODE system describing these two reactions is

$$\begin{cases} \dot{x}_A = -\theta_1 x_A + \theta_2 x_B \\ \dot{x}_B = +\theta_1 x_A - \theta_2 x_B \end{cases} .$$

The Jacobian of the system is

$$\mathbf{J}(\mathbf{x}(t, \boldsymbol{\theta}), \boldsymbol{\theta}) = \begin{bmatrix} -\theta_1 & \theta_2 \\ \theta_1 & -\theta_2 \end{bmatrix},$$

which is singular; hence, the proposed method is not directly applicable.

In this case it is easy to see that $\dot{x}_A + \dot{x}_B = 0$ and the conserved quantity is $x_A(t) + x_B(t)$. The system can be simplified to

$$\dot{x}_A = -(\theta_1 + \theta_2)x_A + \theta_2 x_{\text{total}}, \tag{3}$$

where $x_{\text{total}} = x_A + x_B$. The Jacobian of this one-dimensional system is equal to $-(\theta_1 + \theta_2)$ and does not depend on $x_A$. This system has one non-trivial equilibrium

$$x_A^* = \frac{\theta_2 x_{\text{total}}}{\theta_1 + \theta_2},$$

which is exponentially stable as the only eigenvalue $(-(\theta_1 + \theta_2))$ is negative.

When the system (3) is at steady state, the adjoint state is the solution of

$$\dot{p} = (\theta_1 + \theta_2)p,$$

and is equal to

$$p(t) = e^{(\theta_1+\theta_2)(t-t'')} p(t'').$$

In this case

$$\begin{aligned} &p_{\text{integral}} \\ &= \int_{t'}^{t''} p(s)ds \\ &= \int_{t'}^{t''} e^{(\theta_1+\theta_2)(s-t'')} p(t'')ds \\ &= \frac{1}{\theta_1+\theta_2} p(t'') e^{(\theta_1+\theta_2)(t''-t'')} - \underbrace{\frac{1}{\theta_1+\theta_2} p(t'') e^{(\theta_1+\theta_2)(t'-t'')}}_{=0} \\ &= \frac{p(t'')}{\theta_1+\theta_2}. \end{aligned}$$

The computed $p_{\text{integral}}$ value is used to compute the objective function gradient by S1.

#### Accuracy of gradient computation

The objective function gradient values were computed using ssASA for sensitivities and standard ASA for sensitivities. The values computed with the proposed ssASA method were considered accurate if the error between the two methods was small, with error ($\Delta$) calculated as

$$\Delta = \begin{cases} \left|v^{\text{ssASA}}\right| & v^{\text{ASA}} = 0 \\ \min\left(\left|v^{\text{ssASA}} - v^{\text{ASA}}\right|, \left|\frac{v^{\text{ssASA}} - v^{\text{ASA}}}{v^{\text{ASA}}}\right|\right) & v^{\text{ASA}} \neq 0 \end{cases},$$

where $v^{\text{ASA}}$ and $v^{\text{ssASA}}$ are the gradient values computed with the standard ASA or ssASA method for sensitivities, respectively.

### S3 AMICI implementation

The ASA at steady-state approach was implemented in the AMICI package. AMICI allows for forward integration of differential equation models specified in SBML format or PySB, as well as for forward sensitivity analysis, steady-state sensitivity analysis and ASA for likelihood-based output functions.

For ASA at steady-state, the user can choose between three options:

- only integration, which corresponds to numerical backward integration of the adjoint state ODE, e.g. in the post-equilibration of the ODE

$$\dot{\mathbf{p}}(t, \boldsymbol{\theta}, \mathbf{u}) = -\mathbf{J}(\mathbf{x}(t, \boldsymbol{\theta}, \mathbf{u}), \boldsymbol{\theta}, \mathbf{u})^T \mathbf{p}(t, \boldsymbol{\theta}, \mathbf{u}), \tag{4}$$

  on time intervals $[t'', t_{n_t}), [t_{n_t}, t_{n_t-1}), \ldots, [t_1, t_0)$ with boundary values

$$\mathbf{p}(t_j, \boldsymbol{\theta}, \mathbf{u}) = \lim_{t \to t_j^+} \mathbf{p}(t, \boldsymbol{\theta}, \mathbf{u}) + \sum_{j=1}^{n_y} \left.\frac{\partial h_i}{\partial \mathbf{x}}\right|^T_{(\mathbf{x}(t_j, \boldsymbol{\theta}, \mathbf{u}), \boldsymbol{\theta}, \mathbf{u})} \frac{(\bar{y}_{ij} - y_i(t_j, \boldsymbol{\theta}, \mathbf{u}))}{\sigma_{ij}^2} + \left.\frac{\partial h_i}{\partial \mathbf{x}}\right|^T_{(\mathbf{x}(t_j, \boldsymbol{\theta}, \mathbf{u}), \boldsymbol{\theta}, \mathbf{u})} \frac{(\bar{y}_{i*} - y_i(t'', \boldsymbol{\theta}, \mathbf{u}))}{\sigma_{ij}^2},$$

  and

$$\lim_{t \to t''^+} \mathbf{p}(t, \boldsymbol{\theta}, \mathbf{u}) = 0$$

- only ssASA method, which corresponds to solving the linear system

$$\mathbf{J}(\mathbf{x}^*(\boldsymbol{\theta}, \mathbf{u}), \boldsymbol{\theta}, \mathbf{u})^T \mathbf{p}_{\text{integral}} = -\mathbf{p}(t'', \boldsymbol{\theta}, \mathbf{u})$$

  in the post-equilibration case, or system

$$\mathbf{J}(\mathbf{x}^*(\boldsymbol{\theta}, \mathbf{u}^e), \boldsymbol{\theta}, \mathbf{u}^e)^T \mathbf{p}_{\text{integral}} = -\mathbf{p}(t_0, \boldsymbol{\theta}, \mathbf{u})$$

  in pre-equilibration case.

- combined approach, where ssASA method is attempted first and, in case this fails, numerical integration is used instead.

The alternative approaches are summarized in Figure S1.

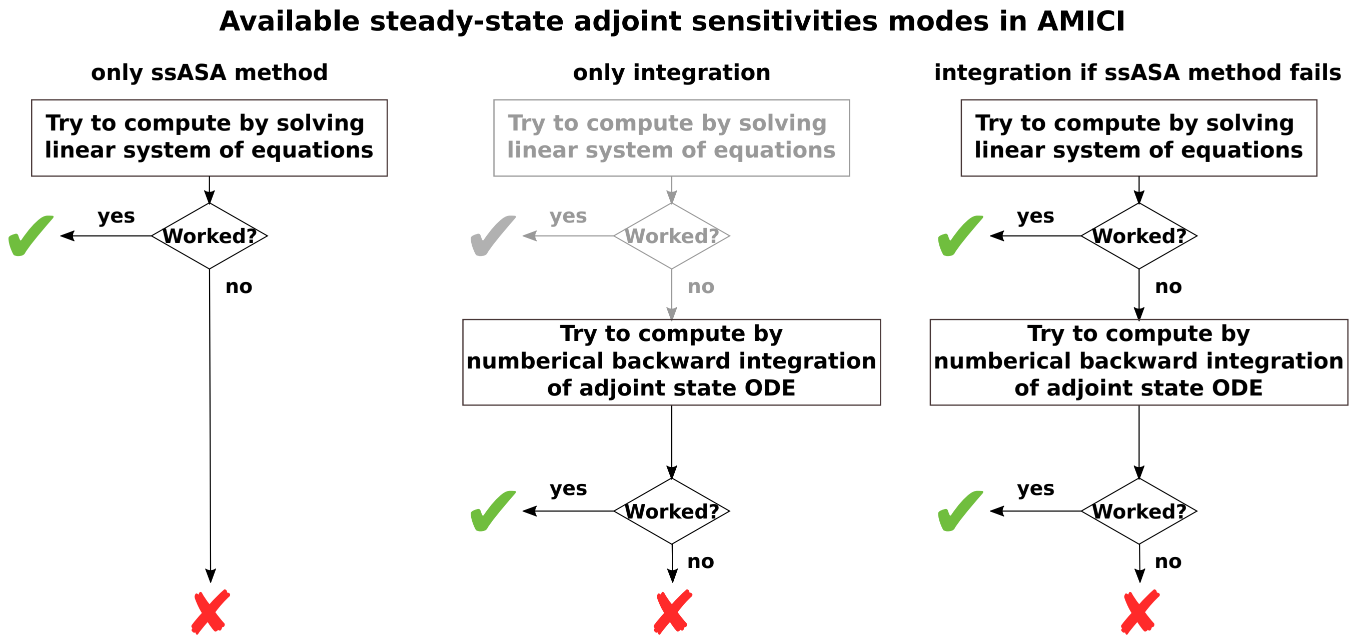


**Fig S1. Available approaches for computing adjoint sensitivities at steady-state in the AMICI package.** For each of the three cases the top box represents the new approach introduced in this study.

### S4 Simulation during optimization with ssASA takes less time

Figure S2 shows the comparison between cumulative simulation time required for optimization using standard ASA (x-axis) or ssASA (y-axis) for sensitivities computation.

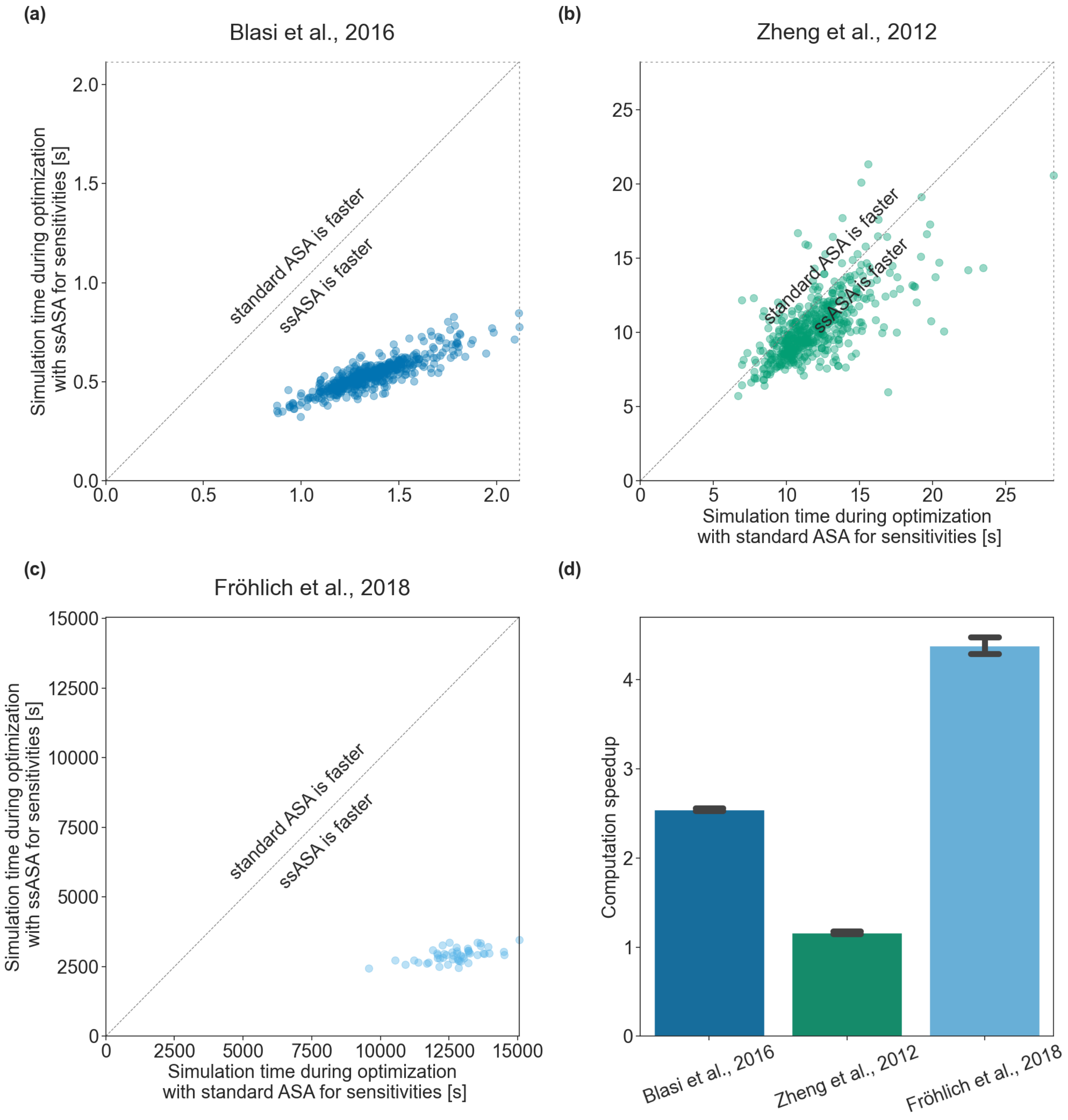


**Fig S2. Cumulative simulation time during optimization.** Each scatter point shows the total simulation time required for one multi-start optimization using standard ASA (x-axis) or ssASA (y-axis) for sensitivities computation. Points on the diagonal correspond to multi-starts that have equal simulation time with both approaches. (a) Blasi *et al.*, 2016 model, (b) Zheng *et al.*, 2012 model (c) Fröhlich *et al.*, 2018 model. (d) Computation speedup of simulation time during optimizations using ssASA for sensitivities computation compared to using standard ASA for sensitivities. Each bar height corresponds to a mean of multi-start local optimization computation speedups and each error bar corresponds to the sample standard deviation.
